# Recombinant Laccase Production Causes Alterations of the *S. cerevisiae* Proteome that are Dependent on the Strain Origins

**DOI:** 10.64898/2025.12.22.696128

**Authors:** Ryan Wei Kwan Wong, Sahil Chandhok, Elizabeth Hui, Thibault Mayor

## Abstract

*Saccharomyces cerevisiae* yeast is a widely used recombinant protein production host. Recombinant protein expression requires adaptation of the host cell proteome to accommodate recombinant expression. However, this adaptation has not been well characterized. A better understanding of the adaptation to recombinant protein expression may inform us of pathways important to the process of expression. The proteome of a laboratory yeast was measured each of the 4 days of recombinant laccase expression to determine the adaptations of the proteome. Whereas a sizeable portion of the proteome had altered levels in response to nutrient depletion in batch growth, a smaller portion of the changes was specific to the laccase expression. By comparing yeast strains of different origins and laccase production capacities, we found that each strain tends to display a distinct response to heterologous expression, regardless of the origin of the laccase. For example, the chaperones Hsp26 and Kar2 were specifically elevated in a whey-derived strain upon laccase expression. Nonetheless, the higher capacity to produce active recombinant laccase in some strains appears to be more strongly associated with small groups of proteins that are constitutively expressed at different levels. These results indicate that strains of different origins each provide a unique cellular milieu that, in some cases, is more favorable for the expression of a given recombinant protein. This study provides potential new targets for strain engineering to improve the yield of select recombinant proteins and provides the first insights into the dynamics of the yeast proteome during recombinant laccase expression.

**Key Points:**

- Proteomes of *S. cerevisiae* strains during recombinant laccase expression determined
- Changes to ribosomal & metabolic protein levels occur during recombinant expression
- Unique cellular milieu, rather than proteome shifts, is linked to higher yields

## Introduction

The baker’s yeast, *Saccharomyces cerevisiae*, has a long history of use for recombinant protein expression dating back to the late 1970s (Beggs 1978; Chevallier and Aigle 1979). A wide variety of recombinant protein products have been produced using *S. cerevisiae* since then, which represents a multi-billion dollar market that includes products such as food additives (Kazemi-Nasab and Shahpiri 2020), industrial enzymes (Kang et al. 2003), vaccines (McAleer et al. 1984) and biopharmaceuticals (Nielsen 2013; Zhao et al. 2024).

Expression of recombinant proteins is taxing on the host cell, requiring diversion of some cellular resources towards producing an exogenous protein. This can potentially create metabolic bottlenecks as cellular resources, such as energy and building blocks are divided between producing the recombinant protein and maintaining essential functions of the host cell (Klein et al. 2015; Zahrl et al. 2019; Kastberg et al. 2022). Additionally, recombinant protein expression can induce significant stress on the host cell from protein folding and degradation burdens (Mattanovich et al. 2004). For secreted proteins these stresses often occur at the ER, engaging the unfolded protein response to manage the stress and triage the protein (Hetz 2012), including degradation via ERAD (Friedlander et al. 2000; Meusser et al. 2005; Ruggiano et al. 2014). For instance, upregulation of the BiP (Kar2) chaperone restored cell growth during recombinant protein expression in *S. cerevisiae* (Kauffman et al. 2002) and BiP is commonly overexpressed to improve recombinant protein yields (Harmsen et al. 1996; Shusta et al. 1998; Kim et al. 2003). Along with the stress response, recombinant expression triggers a transcriptional response in *S. cerevisiae* and other yeast host systems. Transcriptional tuning of stress response, such as the ER chaperone genes *KAR2* and *ERO1*, and of metabolic pathways such as downregulation of glycolytic and TCA cycle genes, has been observed during expression of α-amylase in *S. cerevisiae* (Huang et al. 2017). Similar observations have made in *K. phaffii* expressing multiple different recombinant proteins (Gasser et al. 2007; Yu et al. 2017; Zhang et al. 2020). Transcriptional tuning of cellular metabolism and chaperones in these host cells are a response to nutrient depletion and protein folding stresses during recombinant protein production (Gasser et al. 2007; Yu et al. 2017; Huang et al. 2017). Numerous studies have worked to reduce the impacts of metabolic bottlenecks by altering the growth medium or feeding regiment (Mendoza-Vega et al. 1994; Klein et al. 2015), and to reduce the burdens of recombinant protein expression by overexpressing chaperones to aid in folding (Robinson et al. 1994; Hayano et al. 1995; Shusta et al. 1998) or deleting proteases to reduce degradation (Jones 1991). However, the impacts of recombinant protein expression on the proteome has not been fully characterized, potentially limiting the identification of novel targets to enhance recombinant expression.

*S. cerevisiae* strains from different origins may be better suited to produce specific proteins. Using a large-scale screen, we previously identified subsets of non-laboratory strains that have improved capacity to produce recombinant laccase in comparison to the laboratory strain BY4741 (Wong et al. 2025). These strains are depleted for several genes involved in sugar metabolism such as *HXT* and *IMA* genes, and genes involved in vacuolar degradation (*ATG15*, *CPS1* and *YPS3*). Deletions of some of these genes in the laboratory strain increased production of the heterologous *Trametes trogii* laccase, providing a novel approach to identify bioengineering targets in the laboratory strain BY4741. Interestingly, proteomic analysis revealed that there are also large changes at the protein level between these non-laboratory strains. However, it was difficult to impart which of these changes potentially contributed to higher yield of recombinant production. Therefore, a better understanding of the changes at the proteome level during recombinant laccase expression may help identify proteins key to improving laccase production and heterologous expression.

In this study, we aimed to characterize changes at the proteome level caused by the recombinant expression of a laccase enzyme in the commonly used laboratory strain BY4741. We then determined how these changes vary upon expression of a different recombinant enzyme and in non-laboratory strains with varying abilities to express different laccases. We performed several comparative analyses and identified pathways enriched among altered proteins. Our findings suggest that strains from different origins display distinct response to heterologous expression and the higher capacity to express active recombinant enzymes in some of these strains may be related to unique subsets of differentially expressed in a constitutive manner.

## Materials and methods

### Plasmids and yeast strains utilized

All plasmids and yeast strains utilized in this study are listed in Supplemental File 1. The plasmids were previously constructed in (Wong et al. 2025). The BY4741 strain was a gift from Dr. Phil Hieter (Brachmann et al. 1998). The strains CLIB549, CLIB631 and CLIB649 were a gift from Dr. Joseph Schacherer (Peter et al. 2018).

### Yeast transformation and growth

All growth steps were conducted at 30°C. Yeasts were transformed with either the empty vector (BPM1743), the *T. trogii* laccase *(ttLCC1)* vector (BPM1747) or the *M. thermophila* laccase *(mtLCC1)* vector (BPM1752) using the lithium-acetate/single-stranded DNA method (Gietz and Schiestl 2007), and were grown for 3 days on to yeast extract, peptone and dextrose (YPD) + G418 (200 µg/mL) selective plates.

For experimental cultures, each of the 4 biological replicates contained 10 individual colonies pooled together. An overnight (16 h) pre-culture was grown in YPD + G418 (200 µg/mL), and then the OD_600_ was measured. For the time-course experiments, to maximize consistency within biological replicates for each day, each of the 4 biological replicates started from a master starting inoculum that was split into 4 equal aliquots to be harvested each of the 4 days, for a total of 16 cultures per condition (empty vector & *T. trogii* laccase). 6 mL of each experimental culture was inoculated to a starting OD_600_ of 0.2 (1 mL used for measurement), then four 1 mL aliquots were transferred to a 2 mL 96-deep-well round bottom plate. Each day, one of the 4 aliquots of each strain and replicate was collected into a 1.5 mL microcentrifuge tube, then the cells were pelleted by centrifugation at 3,200 rcf for 5 min. The cell pellets were washed 3 times with 1× Tris-buffered saline (pH 7.4). 100 µL of the cleared media containing the secreted laccase was transferred to a clear 96-well flat bottom plate to be assayed for laccase activity. The cells were then snap-frozen in liquid nitrogen and stored at −70°C for mass spectrometry experiments.

For the multi-strain experiments, experimental cultures were set up as previously described except 3 mL of each experimental culture was inoculated to a starting OD_600_ of 0.2 and 1 mL was transferred to a 2 mL 96-deep-well round bottom plate. The experimental cultures were grown for 4 days (96 h) with shaking at 900 rpm. After 4 days, 100 µL of the cleared media containing the secreted laccase was transferred to a clear-96-well flat bottom plate to be used for laccase activity assays. The remainder of the cleared media was discarded and then the cell pellets were washed and frozen as described previously.

### Laccase activity assay

100 µL of cleared media from laccase expression cultures was assayed for laccase activity. Immediately before quantitation, 100 µL of 2 mM 2,2’-azino-di-(3-ethyl-benzthiazoline-6-sulphonic acid) (ABTS) in 100 mM Britton and Robinson buffer (100 mM each of boric, phosphoric and acetic acid, brought to pH 4 by addition of NaOH) was added to the cleared media to begin the reaction (Alcalde and Bulter 2003). The laccase activity was monitored by UV-visual spectrometry at 420 nm (absorbance, Ab_420nm_) for 1 hour, with readings every minute starting from minute 0, in a Clariostar+ plate reader (BMG). We used shaking between reads in a double orbital shaking pattern at 300 rpm for 30s, followed by centre point reading with 20 flashes. The Ab_420nm_ was plotted against time, and linear regression curves were fitted to the data. To assess only the linear range, data points at the tail end of the curve were trimmed until an R^2^ of at least 0.999 was achieved. Ab_420nm_ was converted to concentration of oxidized ABTS in µmols using the Beer-Lambert law and a molar extinction coefficient of 36,000 M^−1^ cm^−1^ (Childs and Bardsley 1975). Using this, we calculated laccase activity following the definition of 1 unit of laccase activity (1 unit of laccase activity = 1 µmol oxidized ABTS/min).

### Mass spectrometry

Pelleted cells from previously collected cultures were thawed on ice before being lysed in 50 mM TRIS-HCl (pH 6.8) with 2% SDS using a Precellys 24 (Bertin) at 6,800 Hz (3×30s cycles with 90s interval between) and 400 µm acid washed silica beads (OPS Diagnostics BAWG 400-200-04). Cell debris were spun down at 3,220 rcf for 5 min, then the protein concentrations of the cleared lysates were measured using the BCA assay (Thermo Scientific 23225). We digested 5 µg of proteins from the cleared lysates using the SP3 protocol (Hughes et al. 2019), with Sera-Mag™ carboxylate-modified SpeedBeads [E7] and [E3] (Cytiva 45152105050250 and 65152105050250) and sequencing grade modified trypsin (Promega V51135). The digested peptides were acidified with trifluoroacetic acid (TFA) before desalting by StAGE tipping using an AssayMAP Bravo (Agilent) liquid handler and AssayMAP 5 µL C18 cartridges (Agilent 5190-6532).

For STAGE tipping, cartridges were primed with 100 µL of priming buffer (0.1% TFA, 80% acetonitrile (ACN)) then washed with 50 µL of buffer A (0.1% TFA, 5% ACN) before loading of the peptides. A “cup wash” step was performed with 25 µL of buffer A, then a sample wash was done with 50 µL of buffer A. The peptides were eluted from the C18 with buffer B (0.1% TFA, 40% ACN) before drying with a Vacufuge plus (Eppendorf 022820109). Dried peptides were reconstituted in 0.1% FA & 0.5% ACN before loading on the mass spectrometer.

For the BY4741 time-course experiments, an Orbitrap Exploris 480 operated in data independent acquisition (DIA) mode, coupled to an Easy n-LC 1200 (Thermo Scientific) with the temperature set at 7°C was used. 100 ng of each sample was loaded on an Aurora Series Gen3 analytical column, (25 cm x 75 µm 1.6 µm C18; Ion Opticks). The analytical column was heated to 40°C using an integrated column oven (PRSO-V2, Sonation). For this run, buffer A consisted of 0.1% aqueous formic acid (FA) and 2% ACN in water, and buffer B consisted of 0.1% FA and 80% ACN in water. A 60 min gradient was run for the liquid chromatography (LC), beginning with 1 min equilibration at 2% B, then proceeded from 2% B to 20% B over 45 min. The gradient then changed to 32% B over the next 15 min, then to 50% B from 61 to 66 min. Finally, the gradient increased to 95% B over the next 5 min, then held at 95% B for 8 min, before dropping to 3% B over 2 min, and being held at 3% B for 6 min. The analysis was performed at 0.25 µL/min flow rate, with an ion source voltage of 1,900 and the HCD collision energy set to 28%. For acquisition, the initial scan ranged from 380 to 985 m/z and the DIA acquisition fragmented precursors from 379.5 to 980.5 m/z in 10 m/z mass windows overlapping the previous mass window by 1 m/z, with a 2.45 second cycle time.

For the multi-strain experiments, a timsTOF Pro 2 operated in DIA-PASEF mode, coupled to a NanoElute 2 UHPLC system (Bruker Daltonics) was used. 50 ng of each sample was loaded on an Aurora Series Gen3 analytical column, heated to 50°C, and equilibrated with buffer A (0.1% FA and 0.5% ACN in water). The analytical column was then subjected to a 30-min gradient with a 0.3 µL/min flow. Over the first 15 min, buffer B (0.1% FA, 0.5% water in ACN) was increased from 2 to 12%, then from 15 to 30 min it was increased to 33%, followed by increasing to 95% over 30 s and held at 95% for 7.72 min. The DIA acquisition scans ranged from 100 to 1,700 m/z. Following the MS1 scan, 17 PASEF scans of 22, 35 m/z windows ranging from 319.5 to 1,089.5 m/z were performed. Ion mobilities ranged from 0.7 to 1.35 V s/cm^2^ with a 100 ms ramp and accumulation time, and a 9.42 Hz ramp rate resulting in a 1.91 s cycle time. Collision energy was increased linearly as a function of ion mobility from 27 eV at 1/k0 = 0.7 V s/cm^2^ to 55 eV at 1/k0 = 1.35 V s/cm^2^.

Data files from both the Exploris 480 and Bruker timsTOF Pro2 were searched using Spectronaut’s directDIA pipline (Biognosys ver. 19.9.250422.62635), quantifying proteins using the top 3 peptides at the MS2 level. The mass spectrometry files are available on MassIVE (MSV000099162). Data quality was assessed by calculating the coefficients of variation (CVs) and correlation of the data sets using a custom script fitting each data set to a linear model using the “lm” function of the “stats” package in R (ver. 4.5.1). Median normalized protein quantities are listed in Supplemental File 2.

### Principal component and limma analysis

Principal component analysis was conducted using the “PCA” function of the “FactoMineR” package in R and the normalized protein intensities. Limma analysis was performed on median normalized data using the “limma” R package (Ritchie et al. 2015) and a custom script to identify differentially expressed proteins (DEPs). The criteria for hit proteins from limma were as follows: adjusted p-value < 0.05 and log_2_ fold-change < −1 or > 1. P-values are available in Supplemental File 3. Venn diagrams of DEPs were generated using the molbiotools – Multiple List Comparator (https://molbiotools.com/listcompare.php) and edited in Adobe Illustrator.

### KEGG and GO term analysis

Clustered analysis was performed on proteins by plotting their log_2_ fold-change using the “pheatmap” package in R and clustering using the Manhattan distance method. Proteins from each cluster were fed into the ShinyGO 0.85 webtool (https://bioinformatics.sdstate.edu/go/) (Ge et al. 2020) for Kyoto encyclopedia of genes and genomes (KEGG) and gene ontology (GO) term enrichment analysis was performed using ShinyGO 0.85, with a false-discovery rate (FDR) cut-off of 0.05. Significant terms were exported to Supplemental File 4. The Sankey plot was generated using the “sankeyNetwork” function from the “networkD3” package in R.

### Z-score heatmaps

Z-score heatmaps were generated by calculating the z-scores of each protein from the biological replicates of each sample and plotted using the “pheatmap” package in R with clustering using the Manhattan distance method. KEGG and GO term analysis was performed as above.

### Phylogenetic analysis

The “1011distanceMatrixBasedOnSNPs.tab” dataset from (Peter et al. 2018) was used to generate a neighbour-joining phylogenetic tree using the “ape” package in R and exported in Newick format. Genetic distances are calculated as the number of non-identical bases per 100 bases (i.e., percent of non-identical bases). The tree was visualized as a rectangular tree using the Interactive Tree of Life (iTOL) webtool (Letunic and Bork 2024) and was rooted to the strain HN6 from the suspected origin country of *S. cerevisiae*, China (Peter et al. 2018; Duan et al. 2018). The tree was cropped, and strains were highlighted using Adobe Illustrator.

## Results

### Dynamics of the proteome during laccase expression

We sought to characterize the changes at the proteome level in response to recombinant protein expression in yeast host cells over the course of a batch production. We first focused on the expression of a recombinant *Trametes trogii* laccase (TL) in the S288C strain background (BY4741) over 4 days (Fig. 1A), when the heterologous protein reaches its maximum activity (Wong et al. 2025). The laccase was expressed from an episomal plasmid and control cells carried an empty vector (i.e., non-expressing). We measured laccase activity each day to monitor laccase production throughout the course of expression. Laccase activity increased sharply between day 1 and 2, then roughly doubled each day from day 2 to 4 (Fig. 1B). Trypsin-digested cell lysates were run on the mass spectrometer using data independent analysis (DIA). A total of 4,282 proteins were quantified ranging from 4,088 to 4,258 proteins per sample (Supplemental File 2). The quality of the data and reproducibility among replicates was assessed by correlation, with all datasets having a Pearson correlation coefficient of *r* > 0.85, increasing to *r* > 0.98 when comparing replicates from the same day and condition (Fig. 1C). These results suggest that our platform allows good reproducibility and that relatively minor changes occur upon expression of a recombinant protein. Correspondingly, median coefficients of variation (CVs) for each dataset ranged from 6.78 to 11.42% (Fig. S1).

**Fig. 1.**
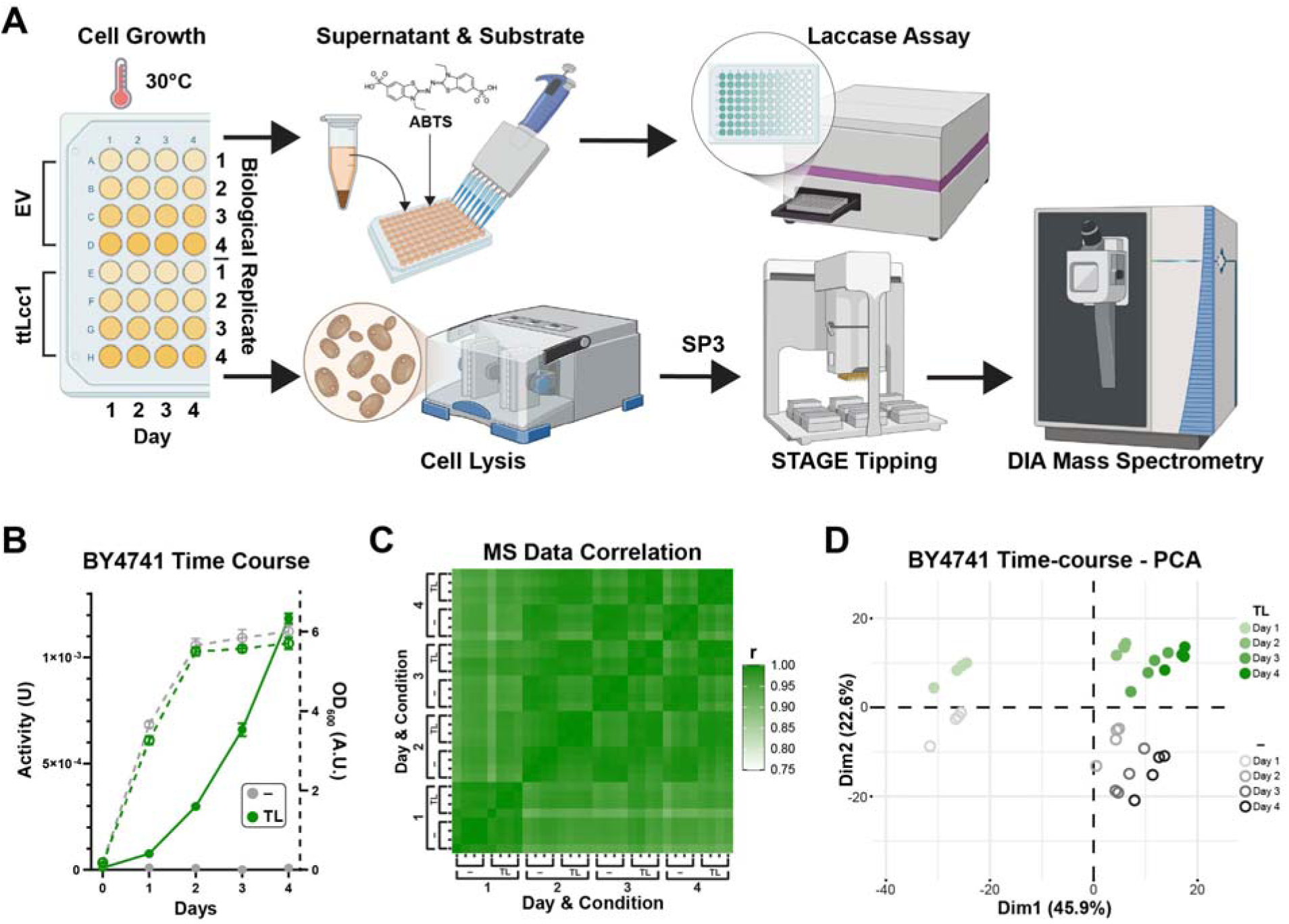
Comparison of the proteomes of BY4741 with and without *T. trogii* laccase expression over 4 days. **(A)** Workflow of the 4-day time-course laccase expression experiment. **(B)** *T. trogii* laccase (TL) activity (Units, solid) and OD_600_ (Arbitrary units, dotted) of BY4741 over 4-days, TL expressing cells in green, and cells with the empty vector (–) in grey. **(C)** Heatmap representing the Pearson correlation of measured protein intensities from each day, condition and replicate. **(D)** Principal component analysis plot of the proteomes of each day, condition and replicate.

To better compare our results, we performed principal component analysis (PCA). The day 1 samples clustered relatively close together in the PCA, with progressively better separation in dimension 1 for samples from each additional day of growth, away from day 1 samples. There was also a stark separation of the samples in dimension 2, bifurcating the samples expressing laccase and those that are not (Fig. 1D). These results suggest that growth, especially during the first day, is most impactful on the changes at the proteome level, while expression of the recombinant protein also clearly contributes to the separation, albeit to a lower extent.

### Metabolic changes during recombinant expression

To specifically delve into the proteomic changes upon laccase expression, we first utilized the “Linear Models for Microarray” (limma) method to identify differentially expressed proteins from our mass spectrometry. Although the limma differential expression analysis method was originally designed for RNA-sequencing and microarrays (Smyth 2004; Ritchie et al. 2015), it has since been frequently utilized for analysis of proteomics data (Schwämmle et al. 2013; van Ooijen et al. 2018; Dowell et al. 2021; Peng et al. 2024) and provides greater quantitative fidelity with low replicates (Dowell et al. 2021). Comparing to day 1, there was up to 631 differentially expressed proteins (DEPs) from a given condition (Fig. 2A, 236 downregulated and 395 upregulated at day 4 when expressing laccase, Supplemental File 3), representing approximately 10.4% of the *S. cerevisiae* proteome altering their levels over the course of the 4 days. Overall, there were more DEPs as time progressed and more proteins with increased abundance compared to decreased abundance. This suggests that upregulation of proteins in response to the demands of recombinant laccase expression is more common than protein down regulation.

**Fig. 2.**
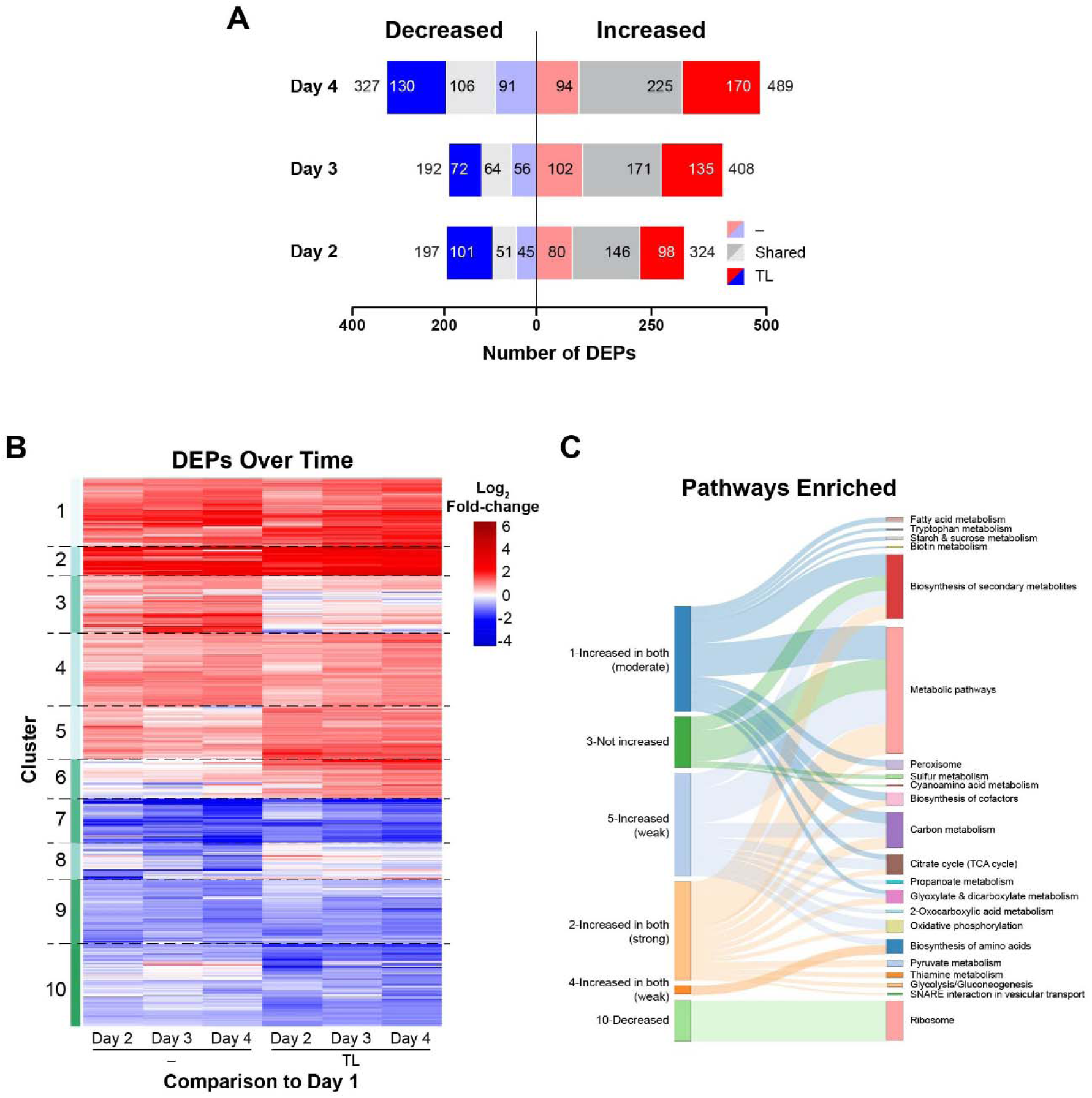
BY4741 time-course comparisons to day 1. **(A)** Bar plot of the number of differentially expressed proteins (DEPs) compared to day 1 that are shared or not between cells expressing *T. trogii* laccase (TL) or carrying an empty vector (–). **(B)** Fold-change heat map of DEPs in comparison to day 1 and clustered using the Manhattan method. **(C)** Sankey plot linking the protein clusters from the heatmap (B) to enriched KEGG terms. Some clusters were not associated to any enriched term. Unless specified, the cluster description is relative to TL expressing cells.

To highlight trends, we clustered the DEPs based on their change relative to day 1. Several clusters encompass a large subset of proteins that increase over time regardless of laccase expression (Fig. 2B; clusters 1, 2 & 4), along with proteins that, in contrast, decrease (clusters 7, 9 & 10, Supplemental File 2). A total of 461 unique proteins are specifically increased or decreased upon laccase expression (clusters 5, 6 & 8, respectively) or not altered when expressing the recombinant enzyme (clusters 3 & 8). To gain a better insight of the functions of the proteins in each cluster, we performed Kyoto Encyclopedia of Genes and Genomes (KEGG) term analysis to highlight the pathways affected. From this analysis, metabolic pathways such as for the biosynthesis of secondary metabolites and amino acids are the most widely shared among proteins expressed at higher levels over time (Fig. 2C). While no KEGG term was associated to individual clusters encompassing proteins reduced over time in both conditions, gene ontology (GO) term analysis of these proteins reveals a conserved reduction of transmembrane proteins involved in carbohydrate transport (Supplemental File 4). These reflect the metabolic changes that *S. cerevisiae* undergoes with nutrient deprivation over the course of the 4 days. Proteins involved in carbon metabolism, citrate cycle (TCA cycle) and oxidative phosphorylation pathways are somewhat more prominently associated to the group of proteins weakly impacted by the laccase expression (Fig. 2C, cluster 5), whereas no pathway is significantly enriched among proteins expressed at higher levels in laccase expressing cells in comparison to the control (cluster 6). Interestingly, the ribosomal proteins are associated with cluster 10, indicating a downregulation of ribosomes during laccase expression.

To potentially better capture the differences caused by laccase expression we repeated the analysis comparing results for each day separately. The total number of DEPs steadily increases over the course of the 4 days (Fig. 3A, Supplemental File 3). This increase is primarily driven by proteins present at higher levels, as the number of proteins with decreased abundances nearly plateaus after day 2, consistent with previous results. Consistently, most differences become apparent following the first day (Fig 3B). As the days elapsed, the divergence of the proteomes becomes more evident (Fig. 3B-F). For instance, the relatively lower levels in methionine/sulfate metabolism proteins (Met3, 5, 10 and 14) is more prominent in days 3 and 4 as these proteins continue to increase over time in absence of laccase expression. Starting on day 2, the hexose transporters Hxt3, 6 and 7 display a marked increase along with the mitochondrial ATP synthase Atp14 and chaperone Hsp10 (Fig. 3D-F). Increased Hxt7 and Hsp10 levels are also major drivers of the laccase expressing proteome signature in the PCA analysis (Fig. S2). The Hap4 transcription factor that promotes the diauxic shift (Blom et al. 2000) was also prominently expressed at higher levels upon laccase expression, which could explain why there was an enrichment of proteins associated to respiration in these cells. We also noted some potential stochastic events such as the high levels of the endosomal membrane protein Vps28 on day 2 (Fig. 3D), which is part of the endosomal sorting complex required for transport-I (ESCRT-I) that plays a role in protein targeting and degradation (Robinson et al. 1988; Rothman et al. 1989; Katzmann et al. 2001). Finally, a glutathione S-transferase (Gtt2) was strongly increased on day 4 (Fig. 3F), possibly to deal with toxins or oxidative stresses induced at later growth stages (Choi et al. 1998). This further highlights the dynamic nature of the *S. cerevisiae* proteome during laccase expression, adapting to the shifting demands of the host cell as nutrients are depleted and more laccase is translated, processed and secreted.

**Fig. 3.**
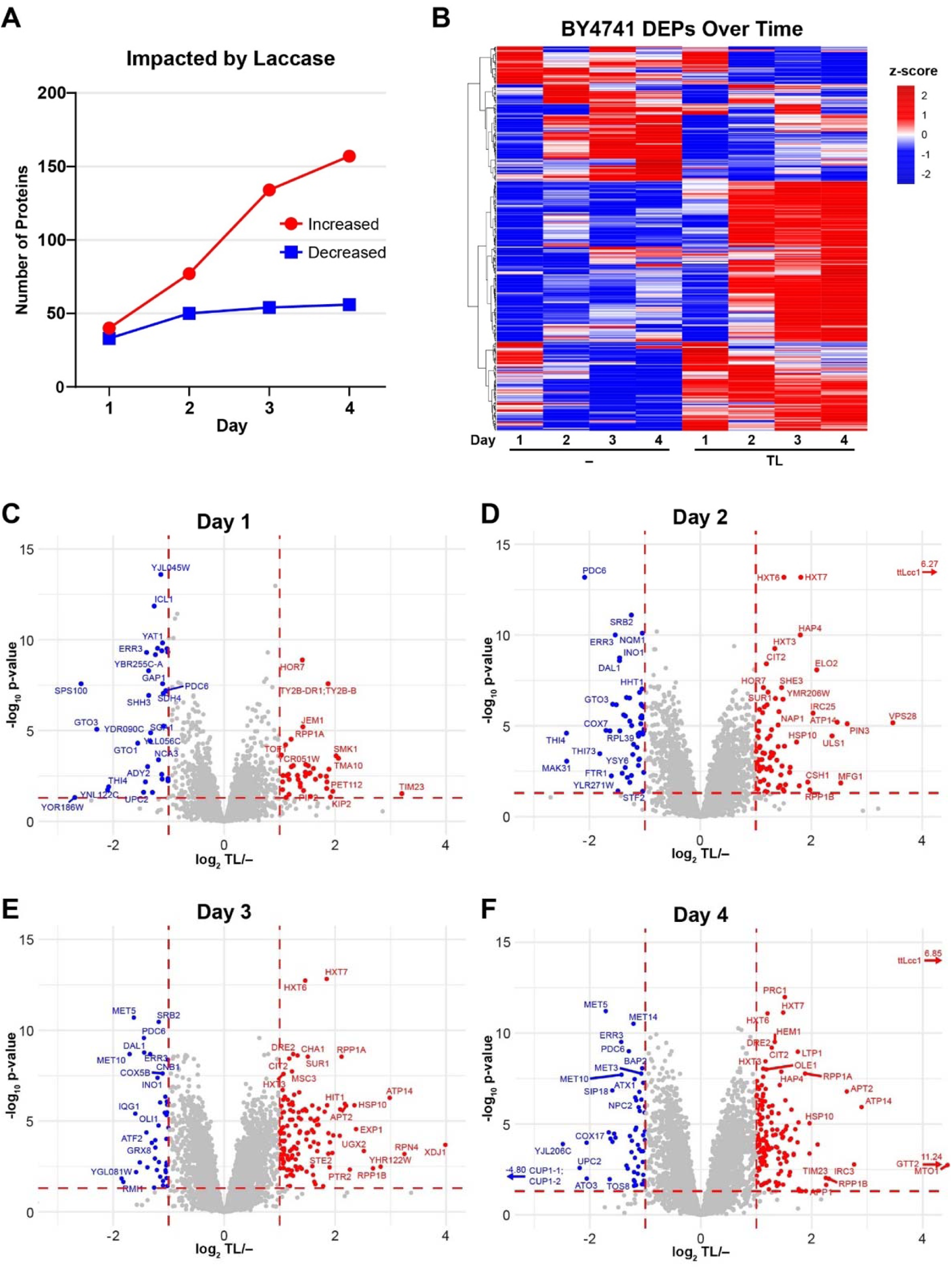
Day-by-day comparison of BY4741 with and without *T. trogii* laccase expression. **(A)** The number of DEPs when comparing BY4741 with *T. trogii* laccase expression (TL) to without (–) each day. **(B)** Z-scores of DEPs when comparing BY4741 with and without TL expression representing proteins present at higher (red) or lower (blue) levels in contrast to other samples. **(C to F)** Volcano plots of the comparisons between laccase expressing and empty vector cells on each day of expression.

### Natural diversity dominates laccase expression-induced proteome changes

When we previously screened a large library of non-laboratory *S. cerevisiae* strains, we identified subsets of strains which better expressed laccases from *T. trogii* and/or *M. thermophila* (Wong et al. 2025). Notably, one group comprised of closely related strains display varied laccase yields, including CLIB649, the top producer of *T. trogii* laccase; CLIB549, a strong producer of *M. thermophila* laccase (ML, which shares only 27.6% sequence identity with *T. trogii* laccase but has a similar protein structure (Wong et al. 2025)); and CLIB631, a strain that did not produce high levels of laccase. The dairy whey-derived CLIB649 strain and cheese-derived CLIB549 strain are more closely related in comparison to the cheese-derived CLIB631 strain (Fig. 4A). For comparison, the W303 strain whose genome virtually identical to S288C from which BY4741 is derived (Brachmann et al. 1998; Ralser et al. 2012) is markedly more distant genetically. We decided to examine the proteomes of these strains more closely to first determine how they are affected by the expression of the two laccase enzymes.

**Fig. 4.**
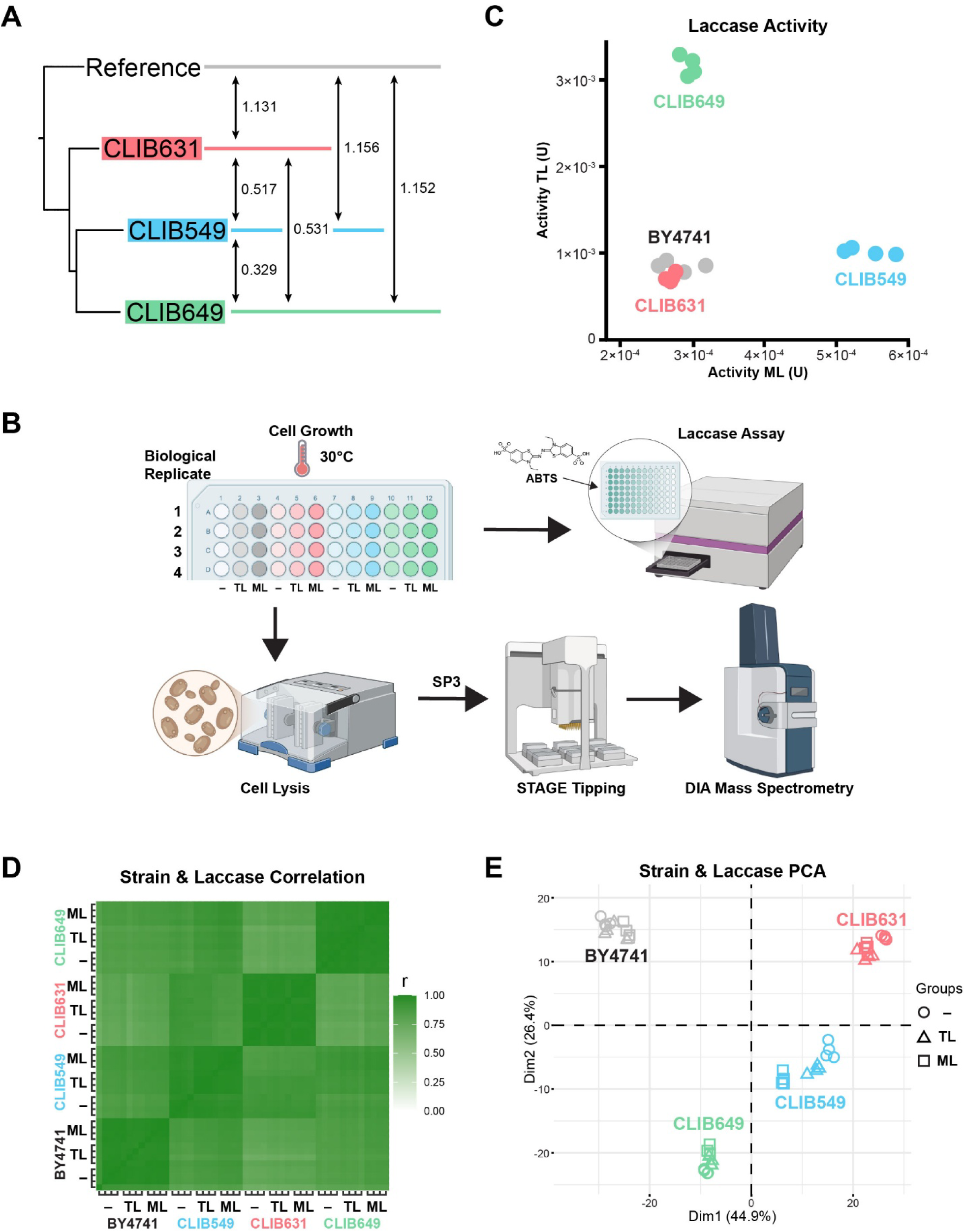
Comparison of the proteomes of related strains expressing different laccases or not. **(A)** Phylogenetic tree of CLIB549 (blue), CLIB631 (red) and CLIB649 (green) in comparison to the W303 reference strain (grey). Genetic distances are displayed between each strain based on SNPs published in (Peter et al. 2018). **(B)** Workflow of the multi-strain experiment with empty vector (–), *T. trogii* laccase (TL) and *M. thermophila* laccase (ML). **(C)** Laccase activity plot comparing *T. trogii* laccase activity (y-axis) and *M. thermophila* laccase activity (x-axis) on day 4 for each indicated strain. **(D)** Correlation heatmap of each proteome from each strain and condition. **(E)** Principal component analysis of the proteomes of each strain and condition.

For each of the four strains (CLIB549, 631, 649 and BY4741) and three vectors (*T. trogii* laccase, *M. thermophila* laccase and empty vector), we grew four biological replicates over four days. The cell-free media from these cultures were tested for laccase activity to quantify the level of secreted laccase expression from each strain. Meanwhile, the cell pellets were lysed for mass spectrometry (Fig. 4B). Laccase activity from CLIB631 was slightly lower than the BY4741 lab strain, CLIB549 had the strongest *M. thermophila* laccase activity, and CLIB649 had the strongest *T. trogii* laccase activity (Fig. 4C), in agreement to our previous results (Wong et al. 2025). We performed data independent analysis (DIA) mass spectrometry on the lysates of these strains, identifying a total of 4,794 unique proteins with an average of 4,617 proteins per strain. Median CVs range between 7.3% to 13.2% (Fig. S3), with high Pearson correlation coefficients (*r*) ranging from 0.91 to 0.998 for replicates (Fig. 4D, Supplemental File 3). Interestingly, each strain displays a distinct signature in the correlation heatmap with stronger correlations within the same strain whether the recombinant laccases were expressed or not (Fig. 4D). To better visualize the effects of the laccase being expressed on each strain’s proteome, we performed PCA on the datasets. There was a strong strain-associated proteomic signature evident in the PCA plots with the data from each strain forming its own distinct cluster in the PCA (Fig. 4E). This indicates that the laccase being expressed does not strongly influence the overall proteome composition of a strain, whereas the strain background strongly dictates the proteome composition. To gain a better appreciation of these differences, we compare the proteome of cells not expressing any laccases between the three genetically similar strains (CLIB549, 631, 649). On average, there was 718 DEPs in these pairwise comparisons (Fig. S4; adjusted p-values < 0.05 and 2-fold or greater change, Supplemental File 3). It was previously shown that protein levels are less variable and more constrained than mRNA abundance between wild strains (Teyssonnière et al. 2024). Nonetheless, a substantial portion of the proteome is expressed at different levels between these strains, which may impact their capacity to produce high yields of recombinant proteins.

### Unique strain response signatures to the expression of laccases

While the laccase being expressed plays a smaller role in defining the proteome when compared to the strain background, it still contributes to specific changes in the proteome. We first compared the impact of the two laccases on the proteome of the different strains. We previously observed that numerous proteins are impacted upon *T. trogii* laccase expression in the lab strain (Fig. 2A), we next wanted to assess whether a similar response can be observed when a different laccase is expressed in that same strain, compared to no expression. From this mass spectrometry run, we identified 34 unique DEPs when expressing either *T. trogii* or *M. thermophila* laccase (Supplemental File 3), and largely the same proteins are up or downregulated in response to either laccase (Fig. 5A). We expanded this analysis to include 135 unique DEPs in any of the four strains (Supplemental File 3). We first observed the levels of these proteins are similarly impacted in a given strain regardless of the expressed recombinant protein (Fig. 5B). These results indicate that the recombinant expression of either laccase causes a similar impact at the proteome level within each strain. An exception to this is a cluster of proteins which show generally higher levels when expressing *T. trogii* laccase in CLIB649, the top producer of that laccase (cluster 3, Supplemental File 2). This cluster showed enrichment of the *protein processing in the ER* KEGG term due to the presence of Emp46, Hsp26, Kar2 (BiP) and Ubc6 in this cluster (Supplemental File 4). Similar to the first proteomic analysis, expression of the recombinant laccases typically triggers more proteins to be expressed at higher levels rather than decreased levels, regardless of the strain (Fig. 5B). Surprisingly, expression of the recombinant laccases causes distinct proteomic signatures in each strain, with mostly unique sets of impacted proteins in each case. These results showcase how seemingly genetically similar strains can respond differently to the stress caused by the recombinant laccase enzymes. These differences may be rooted in their distinct proteome composition and may help explain why different strains are better suited to the production of different recombinant proteins.

**Fig. 5.**
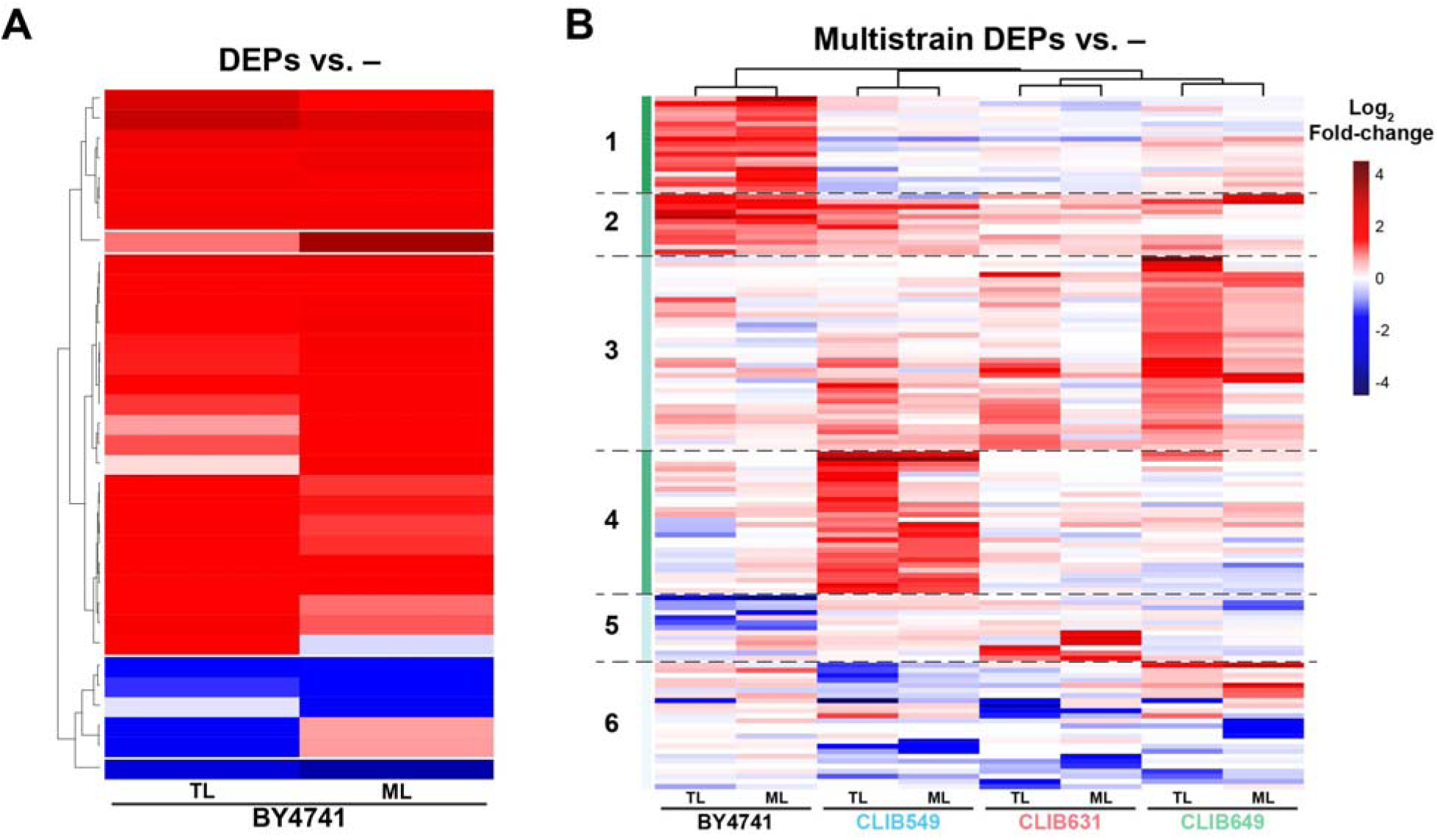
Comparison of the differentially expressed proteins in related strains expressing different laccases. Fold-change heatmaps of proteins in BY4741 **(A)**, as well as in the non-laboratory strains CLIB549, CLIB631 and CLIB649 **(B)**, expressing *T. trogii* (TL) or *M. thermophila* laccase (ML) that are differentially expressed compared to no expression (–) within the same strain. Fold-change scale is shown in (B).

We next sought to determine if any pathways were enriched among affected proteins in the two non-laboratory strains producing high levels of recombinant laccase. Curiously, although CLIB649 is the top producer of *T. trogii* laccase that we have identified, there was no significant KEGG or GO term enrichment found for the proteins increased or decreased when expressing that laccase aside from the previously discussed *protein processing in the ER* enrichment. In contrast, the strong *M. thermophila* laccase producing strain CLIB549, displays enrichment for glucan and carbohydrate metabolic processes, linked to Exg1, Gas3, Pcl10 and Tdh1 that are present at higher levels when expressing the laccase (Supplemental File 4). Nonetheless, these proteins only represent a subset of the impacted proteins in cluster 4. Potentially, there is a lack of concerted response upon heterologous protein production in these cells, or the stress response signatures are masked by higher noise in our analysis.

### Metabolic tuning of the proteome to accommodate expression

We next sought to better understand what proteomic factors might explain how these closely related strains produce different laccases at varying levels to one another. Instead of solely looking at proteins whose levels change in response to recombinant laccase production, we compared the proteomes to identify DEPs that are unique to CLIB649 or CLIB549, when expressing the *T. trogii* or *M. thermophila* laccase, respectively. There are 50 and 24 DEPs unique to CLIB649 and CLIB549, respectively (Fig. 6A & B). Similar to other comparisons, more proteins are present at higher levels. Examination of protein abundances across all samples revealed that these proteins are mostly constitutively expressed at higher or lower levels, regardless of the presence of the recombinant laccase (Fig. 6C & D). Interestingly, DEPs that are always increased in CLIB649 expressing *T. trogii* laccase in comparison to other strains are often involved in fatty acid metabolism (Fox2, Pox1 and Yat1), glyoxylate cycle (Acs1, Ctt1 and Pcs60) and gluconeogenesis (Pdc5, Fbp1) (Supplemental Files 3 & 4), metabolic pathways that work in concert to sustain growth after glucose depletion, indicating a greater reliance on fatty acids as a carbon source in this strain. Accordingly, the fatty acid elongase Elo2 was depleted in CLIB649. However, the transcription factor which controls the β-oxidation, glyoxylate and gluconeogenesis pathways, Asg1 (Jansuriyakul et al. 2016), was decreased in comparison to other strains, perhaps hinting at earlier induction of these pathways (Supplemental File 3). On the other hand, DEPs always increased in CLIB549 and not other strains expressing the *M. thermophila* laccase are chaperone-related (Btn2, Cur1, Ero1, Fes1 and Hrt1) and the cell wall proteins (Css1, Gas3, Utr2, Ynl190w and Zps1), whereas those decreased are involved in thiamine biosynthesis (Pho3, Thi20 and Thi21) (Supplemental Files 3 & 4). In addition to this, comparison of CLIB649 and CLIB549 to the related but poor producing CLIB631 strain highlights a decrease in ribosomes in the previous two strains (Supplemental File 4), mirroring the previous observation in BY4741. These results suggest that the different cellular environment between the strains may enable higher production of distinct recombinant proteins, with possibly an additional contribution from a relatively unique stress response in each strain.

**Fig. 6.**
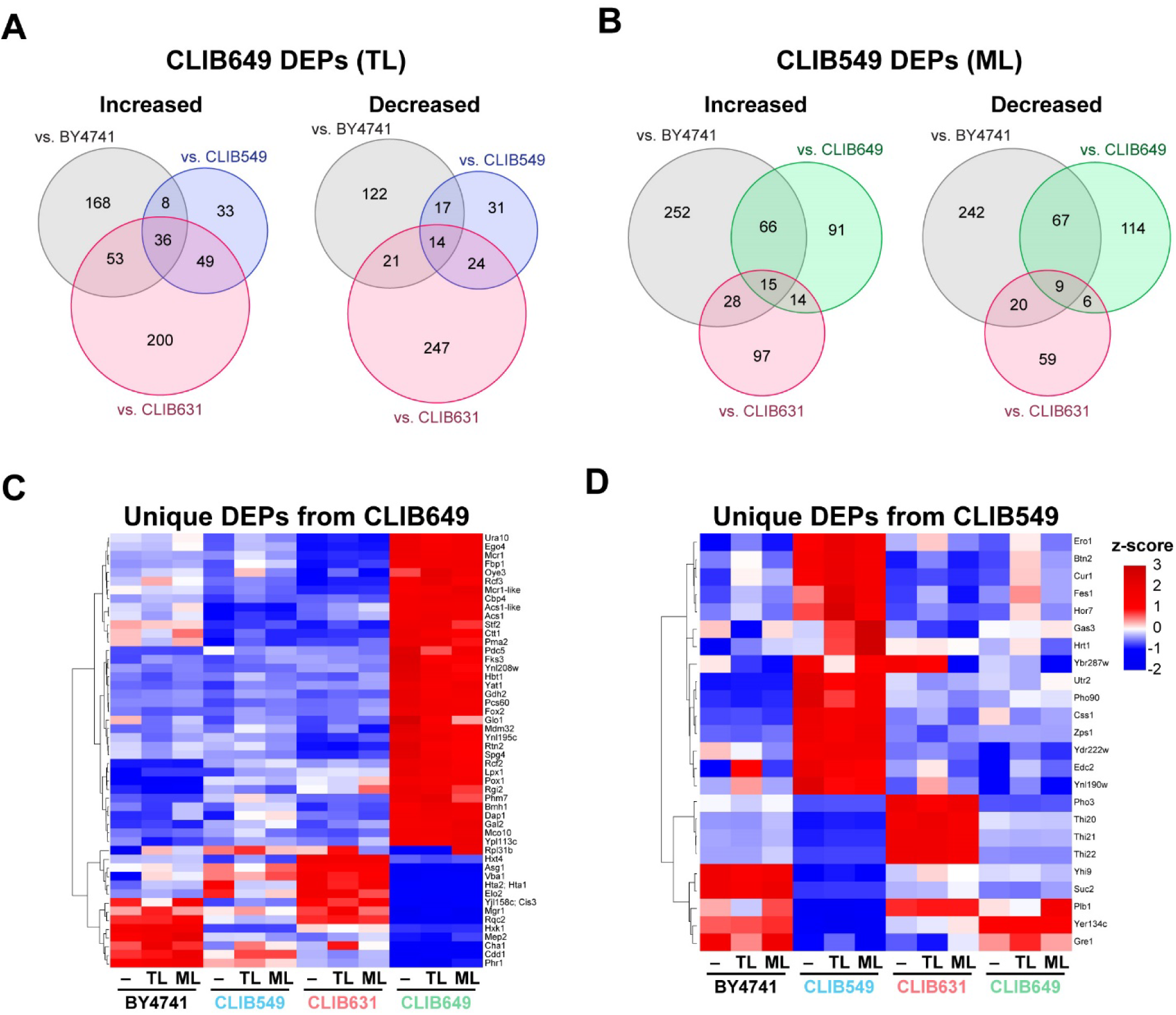
Differentially expressed proteins in top producing strains. DEPs when comparing CLIB649 to the other three strains expressing *T. trogii* laccase (TL) **(A)** and CLIB549 to the other three strains expressing *M. thermophila* laccase (ML) **(B)**. Z-score heatmaps of unique DEPs from CLIB649 expressing TL **(C)** and from CLIB549 expressing ML **(D)** in comparison to other strains expressing the same laccase. Z-score scale as in (D).

## Discussion

Recombinant protein expression is a burden to host cells and requires adaptation to the increased protein load and stress (Mattanovich et al. 2004). While a few proteins, specifically Pdi1 and BiP (which we also observed increased levels under laccase expression), are known to have increased expression levels during recombinant protein expression in yeast (Robinson and Wittrup 1995), large-scale dynamics of the yeast proteome during recombinant expression has not been reported. Specifically, the behaviour of proteins outside of the protein synthesis and secretion pathways during recombinant expression is not well understood. This study provides a direct comparison of several *S. cerevisiae* proteomes during expression of recombinant laccases compared to not expressing, revealing the adaptations of the proteome in response to expressing these enzymes.

Throughout this study we identified several groups of proteins which are specifically and differentially expressed during recombinant laccase expression. We noted that more proteins were upregulated as opposed to downregulated (Fig. 2A, 2B, 3A & 5). These upregulated proteins may potentially be important to sustain high laccase production. However, many studies have instead prioritized gene deletions to assess the effects of certain proteins on recombinant protein expression (Jones 1991; Park et al. 2000; Valkonen et al. 2003; Kim et al. 2015; Strawn et al. 2024; Wong et al. 2025). In the future, it may be important to place greater emphasis on strategies that evaluate how gene overexpression affects recombinant yields.

Several pathways were recurrent in our study, especially when analyzing changes in the commonly used BY4741 lab strain, include glyoxylate cycle, glycolysis/gluconeogenesis, and ribosomes, as well as many proteins involved in transmembrane transport of sugars. Ribosomal proteins were decreased specifically in response to laccase in BY4741, as well as in two strains that produce laccase well (i.e., CLIB549 and CLIB649) in comparison to a similar strain with lower heterologous expression capacity (i.e., CLIB631). Alterations to ribosomal protein levels may itself contribute to improved recombinant protein yield as alterations to ribosomal subunit levels have been demonstrated to improve yields of recombinant Fps1 in *S. cerevisiae* (Bonander et al. 2009). However, the decrease in ribosomal proteins as well as sugar transmembrane transport proteins and increase of proteins involved in glyoxylate cycle and gluconeogenesis are likely primarily part of large-scale proteome re-wiring that is characteristic of diauxic shift regardless of heterologous production, where a change in carbon source decreases the cell’s reliance on sugar uptake and glycolysis (DeRisi et al. 1997; Hiltunen et al. 2003; van Roermund et al. 2003; Stahl et al. 2004; Galdieri et al. 2010; Murphy et al. 2015).

Importantly, the diauxic shift may be important for laccase expression. Notably, we observed laccase production and activity increasing greatly between 24 and 96 hours of expression (Fig. 1B). This timing indicates that the majority of laccase production occurs after the diauxic shift, which is unexpected due to the laccase being expressed under the control of the glucose-dependent *GPD* promoter. Liu, Z. et al. have observed a similar phenomenon when expressing α-amylase (Liu et al. 2012). Much like α-amylase, laccases are relatively complex proteins which require extensive folding and post-translational modifications (PTMs) to reach a highly active conformation (Wong et al. 2025). Two important PTMs for laccases are the addition of copper ions and glycosylation in the Golgi (Tanner and Lehle 1987; Polishchuk and Lutsenko 2013). In agreement with Liu and colleagues, we believe that the transit through the secretory system, as well as the folding and PTMs required to produce active laccase, increases the protein’s dwell time in the cell. This increased dwell time may allow laccase to remain “stockpiled” in the ER until after the diauxic shift occurs, where the conditions for folding may be more favourable (Liu et al. 2012). Similar to this, expanding the capacity of the ER to hold more protein has been effective at improving production of a recombinant protein (Besada-Lombana and Da Silva 2019).

The diauxic shift may create favourable conditions for protein folding, as the shift to utilization of fatty acids and ethanol has a strong requirement for NAD^+^ and NADP^+^, creating NADH and NADPH as a by-product (Hiltunen et al. 2003; Visser et al. 2004). This excess of NADH and NADPH may serve as a reservoir against oxidative stress induced by reactive oxygen species (ROS) created during formation of disulfide bonds during folding (Tu and Weissman 2002), of which *T. trogii* laccase has two(Matera et al. 2008) and *M. thermophila* has three (Ernst et al. 2018). NADH and NADPH may also help to replenish the pool of reduced glutathione, further buffering oxidative stress. Consistent with this, we observed the increase in the glutathione s-transferase, Gtt2, on day 4 of laccase expression in BY4741 (Fig. 3F). Balancing of oxidative stress and folding was observed to improve production of disulfide bond-containing recombinant proteins by Tyo et al. They suggest that when non-stoichiometric ROS and protein folding occur, reduced glutathione is consumed and the cell resorts to formation, breakage and reformation of incorrect disulfide bonds to absorb ROS, creating a futile cycle (Tyo et al. 2012). To balance the folding capacity of the cell, they suggest that downregulation of ribosomes may help to reduce protein translation, thereby matching the amount of folding occurring to the folding capacity of the cell (Tyo et al. 2012). This hypothesis provides a possible explanation for why the reduction of ribosomes is correlated with a slight increase of laccase activity between days 3 and 4 in comparison to days 2 and 3, while the cells biomass marginally increases (Fig. 1B). A similar role for the reprogramming of metabolism to balance redox needs was also observed by Olin-Sandoval et al. where biosynthesis of lysine is downregulated in favour of lysine uptake. This allows NADPH to be used for glutathione metabolism to deal with oxidative stress, rather than for lysine metabolism (Olin-Sandoval et al. 2019). It is perhaps these changes under diauxic shift that shape the cellular milieu and allow certain strains to produce different laccases better than others.

Intriguingly, the alteration of the proteome to recombinant laccase production is mostly unique to each strain, suggesting a strain-specific responses to heterologous laccase expression (Fig. 4E & 5B). Furthermore, each strain responded similarly to recombinant expression of two different laccases (Fig. 4E, 5B, 6C & D), highlighting that the strain background is the strongest determinant of the changes in proteome composition. Nonetheless, the distinct responses within CLIB649 and CLIB549 may also partially account for their higher capacity to produce their respective “preferred” laccase. CLIB649 appeared to adapt its proteome to the changing carbon source needs better than other strains by upregulating fatty acid metabolism, glyoxylate cycle and gluconeogenesis pathways to counter depleting glucose levels. This, perhaps, also helps buffer the oxidative stress of folding *T. trogii* laccase within CLIB649. Meanwhile, CLIB549 may require more direct handing of stress from *M. thermophila* laccase production by upregulating chaperones and related proteins to negotiate the protein folding stress (Supplemental Files 3 & 4). The expression difference of the two laccases could be explained by specific structural constrains of each enzyme despite their conservation (e.g. the *M. thermophila* laccase has three disulfide bonds to potentially sustain higher temperature (Ernst et al. 2018)). Our observations suggest that overexpression of proteins may be under explored for boosting recombinant protein production and that subtle differences in a strain’s proteome and adaption response may affect a strain’s ability to produce a recombinant protein.

## Supporting information

Supplemental File 1 - Yeast and Plasmids Used

Supplemental File 2 - Proteins Quantified

Supplemental File 3 - Statistical Tests

Supplemental File 4 - KEGG and GO Tables

## Acknowledgments

We thank all the members of the Mayor lab for discussions during the course of this project, Yuming Shi for his assistance with data analysis and Jason Rogalski, Renata Moravcová, Jeanne Yuan and Lok Tin Hui for mass spectrometry instrument maintenance and sample loading. Figures 1A and 4B were created with BioRender.

## Author Contributions

R.W. designed, executed and analyzed the data of most experiments. S.C. generated the fold-change and z-score heatmaps. E.H. assisted in preparation of mass spectrometry samples. T.M. helped design the experiment and analyze the data. The manuscript was written by R.W. and T.M. and edited by all the authors.

## Data Availability

All data generated or analyzed in this study are included in this published article and its supplemental material, or available online at MassIVE (MSV000099162). Computational scripts are available at www.github.com/RyanWKW/Recombinant-Laccase-Production-Causes-Alterations-of-the-S.-cerevisiae-Proteome.

## Author Information

Ryan Wei Kwan Wong

Sahil Chandhok

Elizabeth Hui

Thibault Mayor

## Ethics Approval and Consent to Participate

Not applicable.

## Consent to Publish

Not applicable.

