## Supplemental Figures for "Recombinant Laccase Production Causes Alterations of the *S. cerevisiae* Proteome that are Dependent on the Strain Origins"

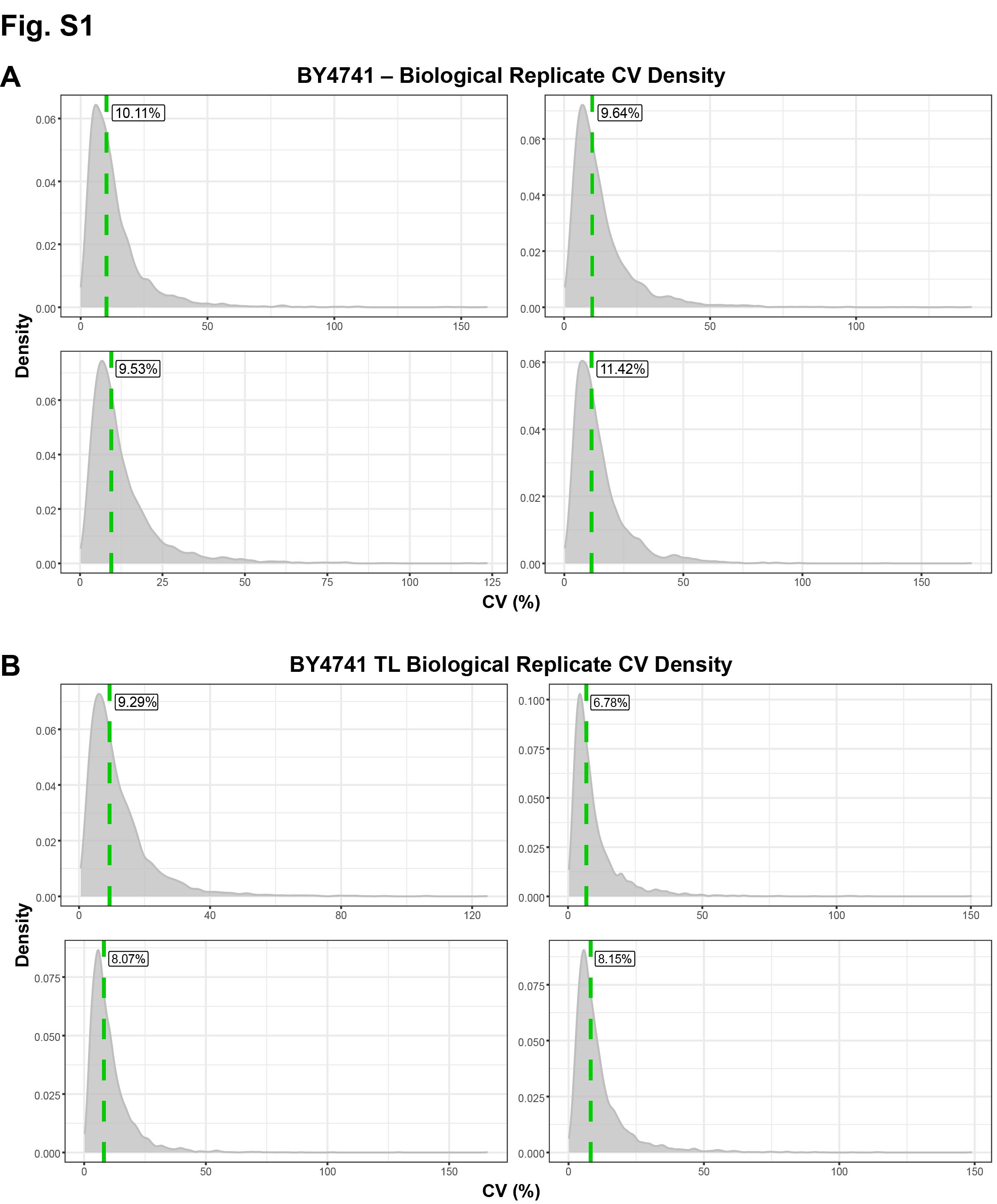

**Fig. S1 Median CVs of BY4741 replicates (A)** CV density plots of the non-expressing condition and **(B)** of the *T. trogii* laccase expressing condition. Median CVs are labeled and indicated by green dashed line.

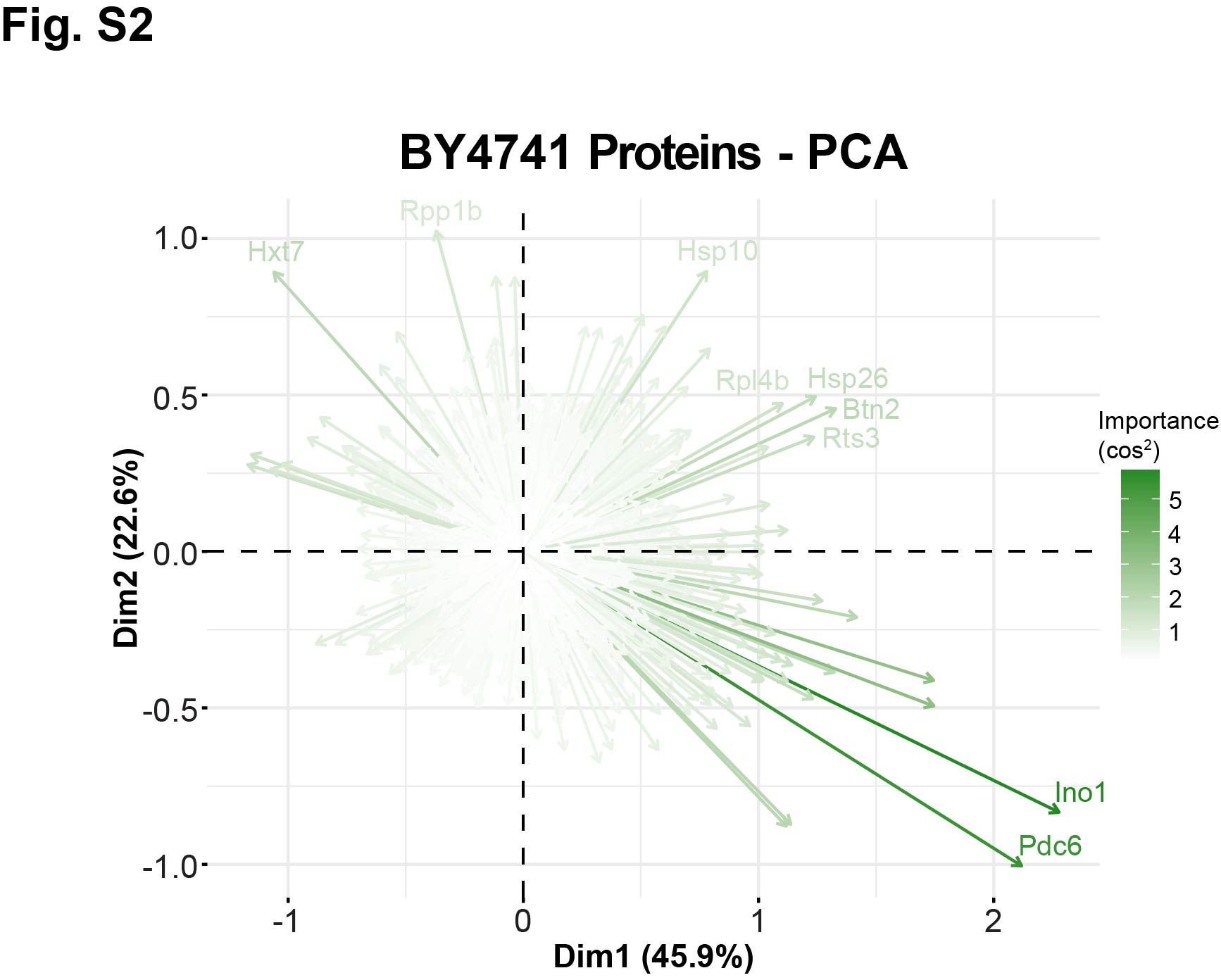

**Fig. S2 Protein contributions to proteome signature** Principal component analysis plot displaying the contributions of each protein towards signatures of older cultures (rightwards) and of *T. trogii* laccase expressing (upwards) or non-expressing (downwards).

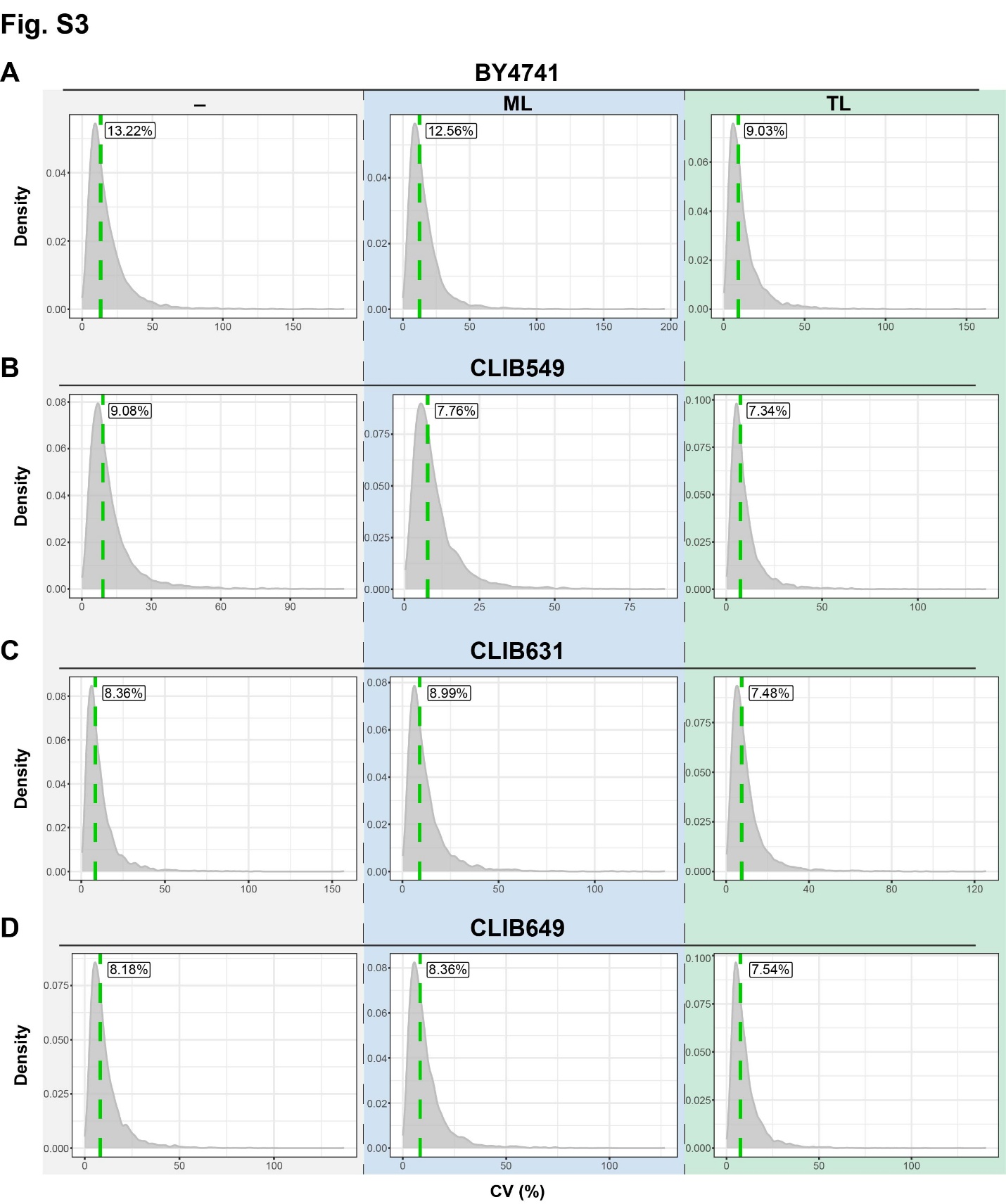

**Fig. S3 Median CVs of each strain and condition** CV density plots of **(A)** BY4741, **(B)** CLIB549, **(C)** CLIB631 and **(D)** CLIB649. Median CVs labeled and indicated by green dashed line.

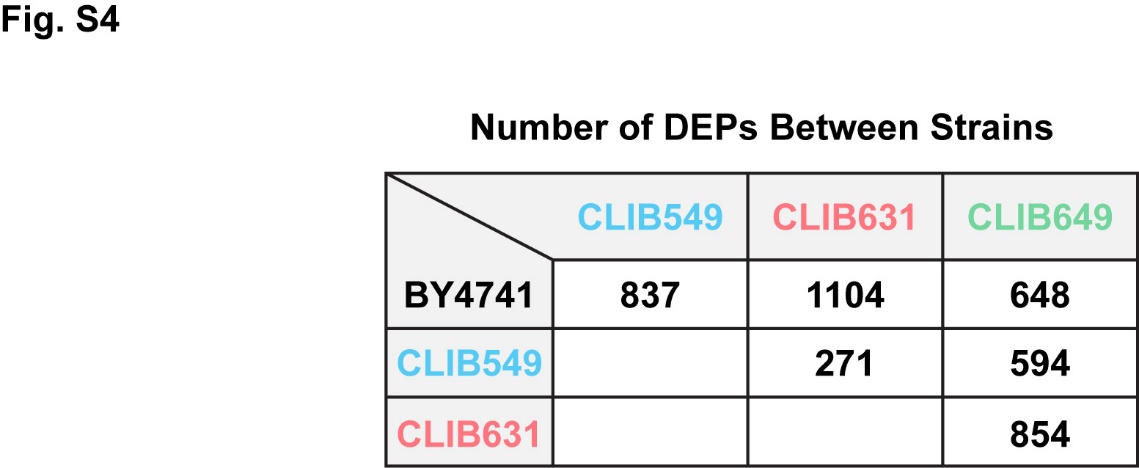

**Fig. S4 Differentially expressed proteins in non-expressing conditions**
